# Engineering Proprioceptive Implants via Surgical and Regenerative Approaches: Preliminary Interpretations

**DOI:** 10.1101/2022.04.03.486912

**Authors:** Siddhartha Das, H.D. Sarma, Jayesh Bellare

## Abstract

The periodontal ligaments are a group of specialized connective tissue fibres with vascular and neural elements that essentially attach a tooth to the alveolar bone. Endosseous dental implant replacing a lost tooth, gets ankylosed to the alveolar bone without intervening periodontal fibres (osseointegration). Hence, proprioception, one of the most important function of periodontal ligament is not elicited by commercial dental implants currently in use. To salvage the flaw, in our proof-of-principle trial in rabbits, biodegradable nanofibres were coiled around the additive manufactured (AM) customized titanium implants. Further, human dental pulp stem cells (DPSCs), adult mesenchymal stem cells of neuro-ectodermal origin, were seeded on the nanofibrous coated, orthotopically placed 3D-printed titanium implants and were induced to differentiate into neural cell lineages. The invivo anchoring mechanism of these biodegradable neuro-supportive scaffold coated implants could probably be “proprioceptive osseointegration” instead of defaults events leading to normal “osseointegration” and could exhibit features similar to periodontium, having possible anastomosis between the severed nerve terminals present in the wall of the extraction socket relaying to/from brain and newly differentiated neural cells present in the regenerated neo-tissue complex, gradually replacing the biodegradable scaffold and may eventually results in the development of proprioceptive osseointegrated root-form endosseous dental implants in near future.

## Introduction

Co-ordinated and precise oral-motor movements depend on afferent inputs from specialized sensory organs in the orofacial region. Among others, a healthy natural tooth consists of extremely sensitive mechanoreceptors located in the periodontal ligament (PDL)^1^ cushioning its root(s). These components largely serve in motor behaviour of maxillo-mandibular complex involving functions such as mastications, biting etc. Earlier studies have established the rich sensory innervation of periodontal ligament via the trigeminal pathways through superior or inferior alveolar branches.^2^ The nerve fibre enters the PDL in the apical region and perforating the lateral wall of the alveolus, give rise to the plexus of nerves. The single myelinated nerve fibres originate from the main nerve bundle lose their myelin sheath and further terminates in free nerve endings, Ruffini like mechanoreceptors, Meissner corpuscles and spindle like pressure/vibration endings.^3^ Activity of these sensory receptors relies on stimulus applied on teeth and supporting structures and transmits afferent information to the central nervous system.^1,4^

In cases of edentulism, prosthodontic treatment considerations such as dental implants are favoured and advocated.^5^ Although oral implant enjoys a high success rate and is secured in bone through a process called osseointegration,^6,7^ the exclusion of PDL and its functionality such as proprioception after the healing events render the conventional osseointegrated implant rather a compromised prosthetic substitute.^8^ Of note, years of research studies lead to the evidence of probability to a widespread and admitted concept of interruption of neural circuitry in osseointegrated dental implants, following absence of the periodontal mechanoreceptors in the juxtaposed tissue, effectuating failure in the propagation of primary sensory inputs.^2,9−12^ Hence, to address the pitfall, numerous contemporary attempts towards the engineering of PDL tissues, interfacing alveolar bone or endosseous implants have been reported globally.^13−23^

The innervation of the aforementioned PDL via the trigeminal nerve for proprioception in humans has been studied extensively.^24^ Briefly, the peripheral part of the trigeminal nerve consists of CN V1 (ophthalmic div.), CN V2 (maxillary div.) and CN V3 (mandibular div.). The purely sensory, CN V2 (maxillary nerve), emerges between the ophthalmic and mandibular nerves, courses and divides into anterior and middle superior alveolar branches further continuing in the anterior and lateral wall of maxillary sinus to innervate anteriors and premolar teeth respectively. Additionally, posterior superior alveolar nerves also arises from CN V2 in the pterygopalatine fossa and continues through the infratemporal surface of the maxilla and subsequently innervates second and third molars, the roots of the maxillary first molar, buccal mucosa and posterior maxillary gingivae.^25^ The CN V3 (mandibular div.), a mixed cranial nerve continues from the lateral part of trigeminal ganglion and is composed of large sensory and small motor root. The sensory root innervates the mandibular teeth and gums, auricle, external acoustic meatus, skin of temporal region, tympanic membrane etc.^26^ while the motor branch innervates the muscles of 1st pharyngeal arch.^27^ The peripheral sensory branches from CN V1, CN V2 and CN V3 gradually merge at the trigeminal ganglion (Gasserian ganglion/semilunar ganglion) located in the depression on the petrous apex of temporal bone known as Meckel’s cave.^28^ Dorsal to the trigeminal ganglion and ventral to the pons, trigeminal root consisting of large sensory and small motor division, leaving the pons at the root exit zone could be noted. The sensory root consisting of central processes of pseudo-unipolar neurons in the trigeminal ganglion, extends into pons and communicates with nuclei in the brainstem. The sensory nucleus extends from the midbrain superiorly to upper cervical spinal cord and consists of mesencephalic, principal and the spinal trigeminal nuclei. On the other hand, the trigeminal motor nucleus is located in the mid-pons, medial to the principal sensory nucleus and could be traced back to its union with CN V3.^29^

The forenamed mesencephalic nucleus, a slender columnar structure, extends in the dorsomedial tegmentum from the level of the trigeminal motor nucleus in the pons to the rostral midbrain^30^ is rather unique for containing cell bodies of primary sensory neurons conveying information from proprioceptors present in oculomotor/masticatory system^31,32^ and are involved in a wide range of activities such as pressure sensation from teeth, palate, TMJ capsule, etc.^33^

The principal sensory nucleus receives general somatic afferent fibres and consists of ventrolateral and dorsomedial nuclei. CN V1, CN V2, and CN V3 supplies information to the ventrolateral nucleus while primary afferent inputs from the oral cavity transduce information to the dorsomedial nucleus.^34^ The extension of spinal nucleus of trigeminal nerve is caudally to the outer lamina of the dorsal horn of the upper three to four cervical spinal segments and contains three different subnuclei in a rostrocaudal direction: the subnucleus oralis, the subnucleus interpolaris, and the subnucleus caudalis.^34^ Subnucleus oralis receives afferent inputs from nasal and oral regions,^35^ subnucleus interpolaris and subnucleus caudalis receives information from the cutaneous portions of the face however the latter receives extensive sensory inputs from the cheek, jaw, and forehead.^34^ Medial to the principal sensory nucleus in the mid-pons lies the motor nucleus and conveys special visceral efferent fibers.^36^ The fibres of the motor nucleus is closely associated with mesencephalic nucleus and in cooperation regulates the bite force.^36,37^

The fibres ascends via various tracts and/or lemniscus for their respective projection in the thalamus [venteroposterior (VP) nucleus], continues through the posterior limb of the internal capsule and eventually allowing the oro-facial area to be represented in the postcentral gyrus, the sensory cortex, or Brodmann areas 3, 1, and 2.^36,38^

In view of the above discussed trigeminal system and its central connections for oral proprioceptive function in humans, we hypothesized that our modifications of coating systems^39−41^ such as neuro-supportive nanofibrous coating with exogenous mesenchymal stem cells^42^ in endosseous dental implants can generate proprioceptive features. We, therefore, decided to investigate the proof-of-principle mechanism in rabbit study models because its trigeminal system^43^ and somatotopic organization^44^ shares similar homology to that in humans.^45−47^ Hence, the customized endosseous titanium dental implants were conceptualized, designed, and later fabricated by additive manufacturing (AM)/manually and were further subjected to post-fabrication electrospun neuro-supportive nanofibrous coating in conjunction with exogenous mesenchymal stem cells for orthotopic implantation in rabbits to reinstate proprioceptive functions.

## Methods

### 0.1. Procurement Of Animals

The proof-of-concept animal trials were conducted at Experimental Facility & Radioisotope Laboratory, Radiation Biology and Health Sciences Division (RBHSD), Bhabha Atomic Research Centre (BARC), Mumbai, under the Department of Atomic Energy, Government of India. The research protocol was reviewed and approved by the panel of the ethics committee for laboratory animal research of BARC, RBHSD, Mumbai, BARC-IITB (Animal study proposal no. BAEC /14/2019), and was performed in accordance with the guidelines of the Committee for the Purpose of Control and Supervision of Experiments on Animals (CPCSEA), Ministry of Social Justice and Empowerment, Government of India. In addition, all experimental surgical protocols and procedures as well as the post-operative care and monitoring were supervised by the senior scientific officer and respective staff of RBHSD, BARC, Mumbai. The methodology for the present study is as per International Standard ISO-10993-677.^48^

Three healthy adult male New Zealand white rabbits were used in the study (Supplementary Tables).The pathogen free rabbits were obtained from Reliance Life Sciences Pvt. Limited, Mumbai, India, after due approval from the IAEC. Each rabbit was numbered and was kept in cages under standardized conditions.^49^ The animals were kept on a 12-hour day/night cycle with *ad libitum* access to food and water.

### 0.2. Radiographic Planning & Measurements

Series of approximate dental measurements were considered in rabbits with Planmeca ProMax® 3D Mid imaging unit (Planmeca, Helsinki, Finland) and saved as DICOM files and were later analysed with dedicated softwares (OsiriX® 8.5, Osirix Foundation, Geneva, Switzerland, https://www.osirix-viewer.com and Planmeca Romexis version 4.6.0, Planmeca, Finland). Mesiodistal width, buccolingual width, mid-point of incisal margin to apex length, a crude estimate of the curvature in mandibular left central incisor and pre-implantation evaluation of jaw anatomy were thoroughly investigated by an experienced bioengineer and surgeon (Supplementary Fig). The range of dimensional values obtained specifying the tooth of interest i.e., mandibular left central incisor, (301, modified Triadan system^50−51^) was utilized for fabricating the implants.

**FIG. 1.**
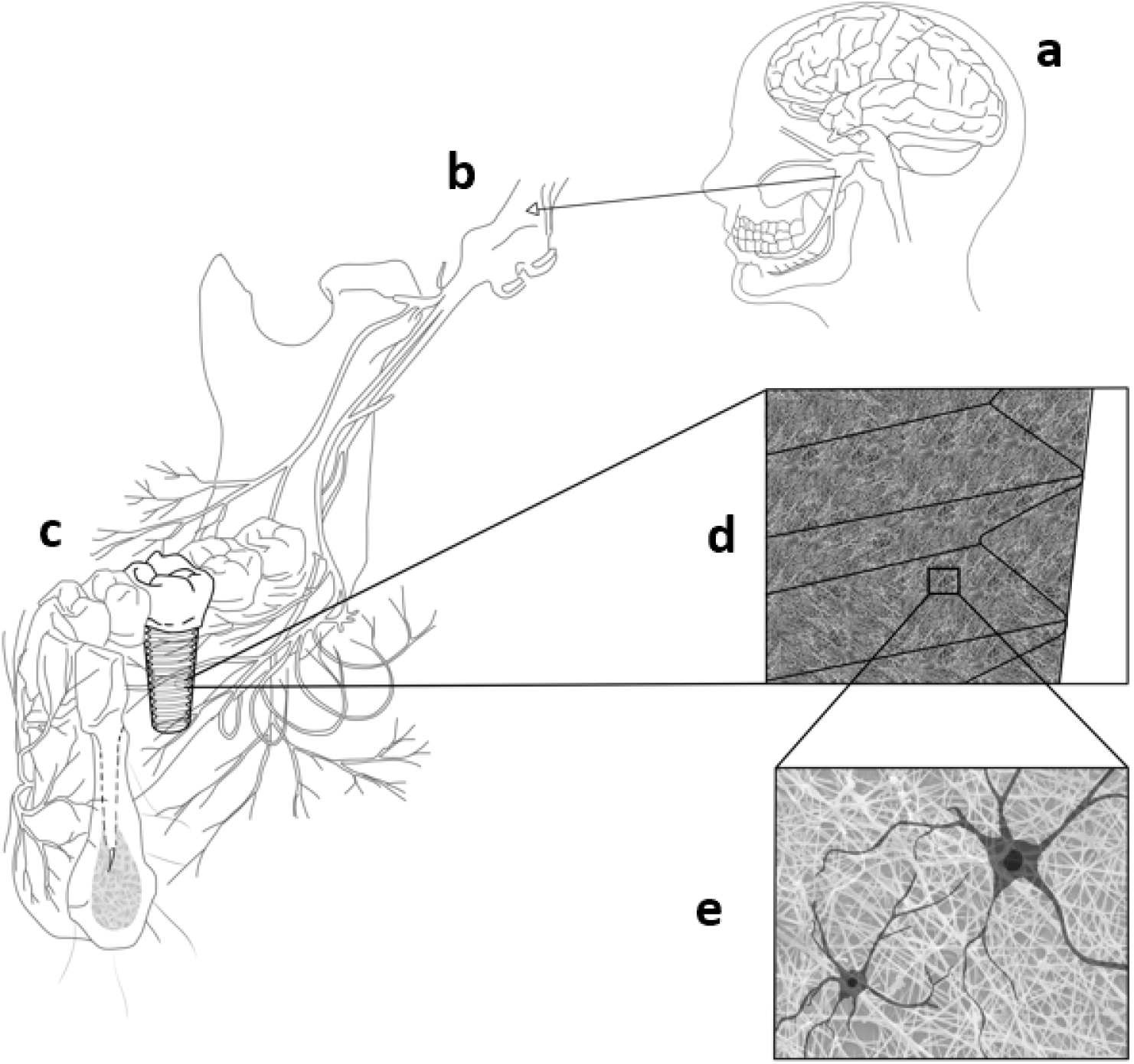
Probable mechanism for the development of proprioception in the modified root-form endosseous dental implant: Here, a portion of the mandible with its neural innervation is depicted for understanding the significance. a - Human brain with trigeminal ganglion and its branches - ophthalmic (V1), maxillary (V2), and mandibular (V3) branches. b - The posterior division of the mandibular nerve gives off sensory branches viz. lingual, long buccal and inferior alveolar nerves. c - Portion of the mandible with anterior and posterior teeth and modified dental implant replacing a posterior teeth, d - Magnified image of a portion of modified dental implant coated with electrospun nanofibres, seeded with exogenous dental pulp mesenchymal stem cells, e- Fraction of mesenchymal stem cells in the nanofibrous coating are differentiated to neural cells upon receiving appropriate signals. These differentiated neural cells present in the regenerated neo-tissue complex may anastomose with the severed/terminal sensory branches of alveolar division originating from trigeminal nerve supplying the wall of the alveolar socket during the healing phase and thereby completing the neural circuitry, resulting in the development of proprioceptive sensation in those implants.

### 0.3. Implant Fabrications

Seven titanium dental implants were shaped and fabricated manually from intramedullary titanium elastic nail (1 *×* 4 mm) (Uma Surgicals, Mumbai, India) in the workshop of MOD Lab, Design and Inspection Section (EmA&ID), BARC, Mumbai, by an experienced bioengineer and surgeon. The macro design features with trapezoidal cross-section (n = 7, diameter = 3.65 ± 0.081 mm, best conformed/suited, n = 1 is installed) for the tooth of interest were in compliance with the precise information acquired through CBCT imaging of the craniofacial region in rabbit study models (Supplementary Fig). Likewise, 3D printed customized rabbit dental implants (n = 7, diameter = 3.65 ± 0.081 mm, best conformed/suited, n = 1 is installed) were designed using the 3D design software, Solidworks 2018 (Dassault Systemes Solidworks Corporation, Waltham, MA, USA) and exported as .STL files by an experienced bioengineer and surgeon (Supplementary Fig). A DMLS additive manufacturing system (EOS M280, EOS GmbH, Krailling, Germany) was used to print the biocompatible titanium (6%)-aluminium-(4%) vanadium extra low interstitial (ELI) grade 23 powder with particle size distribution of 5 - 55 μm. A ytterbium laser system was utilized for layer by layer fabrication of implants with 33 W laser power, 1000 mm/s laser speed and a wavelength of 1060-1100 nm, with layer size of 60 μm (Supplementary Fig). Post-production cleaning steps were followed as described previously.^52^

### 0.4. Stem Cell Culture

Human Dental Pulp Stem Cells were obtained as described previously^42^ and were maintained in low glucose MEM-alpha (Gibco, Invitrogen, Carlsbad, CA, USA), supplemented with 18% fetal bovine serum (MSC-FBS, Gibco) and 100 g/ml streptomycin (Invitrogen/Gibco, Grand Island. NY; US), 100 U/ml penicillin (Invitrogen/Gibco), 0.1 mM ascorbic acid (Sigma-Aldrich Corporation, St. Louis, MO, US), and 2 mM L-glutamine (Gibco). Cells were maintained at 37 °C in a humidified atmosphere of 5% CO_2_.

### 0.5. Coating Of Titanium Implants

The fabricated implants (manually prepared and 3D printed implants) of size 3.65 ± 0.081 mm diameter *×* 27 ± 0.81 mm length were coated with neuro-supportive nanofibres composed of a composite blend of polycaprolactone (Mw-80,000) 8.33 (w/v) % (Sigma Aldrich, USA), gelatine type A 0.833 w/v % (Sigma Aldrich, USA), in 2,2,2-Trifluoroethanol (Sigma Aldrich, USA).^42^ Uncoated titanium rabbit dental implant was used as controls (Supplementary Fig). The technology for fabrication of neuro-supportive nanofibrous coating on the surface of titanium implants by modified electrospinning technique has been described previously by the same author^42^ and was performed by carefully placing the fabricated dental implant (attached to the rotating shaft of DC motor) in between the syringe tip and collector plate^42^ in the *Silicate Lab of J*.*B*., *Chemical Engineering, IIT Bombay* (Supplementary Fig). The modification of the electrospinning apparatus for fabricating nanofibrous coating, its subsequent physico-chemical characterization, and invivo response have been described exhaustively in our earlier and related study.^39−40^ The thickness of the elastomeric porous neuro-supportive nanofibrous coating on the surface of dental implants was chosen carefully to permit proper insertion (i.e. just enough loose to allow inward movement in the fresh extraction socket) and adequate intimate contact of nanofibrous coating with the inner wall of fresh extraction socket for ensuring necessary primary stability for uneventful healing, post-surgery. Prior to surgery, the uncoated and neuro-supportive nanofibrous coated implants were sterilized by irradiating with 25 KGy of gamma exposure in a gamma chamber (GC-900, BRIT India make), Food Technology Division, BARC, Mumbai, India. Samples for gamma irradiation were delivered in sealed and sterile containers.

### 0.6. Stem Cells & Coating

The Human Dental Pulp Stem Cells (hDPSCs) were trypsinized after reaching a state of 80% confluency (3*×*10^6^ cells/flask/vial) and were suspended and agitated gently in vials (n = 2) with 2 ml complete media to generate a homogeneous solution. Sterile insulin syringes (BD 1-mL conventional insulin syringes) was used to insert homogenous cell suspension inside the layers of nanofibrous coating by an experienced bioengineer and surgeon. Additionally, the cell suspension (vial/implant) was pipetted carefully on the sterile surface of the nanofibrous coated implants whilst applying gradual rotational motion along its transverse axis. The nanofibrous coated implants with stem cells were incubated for 15 mins and were transferred into a 24 well cell culture plate and further incubated in complete medium under conditions of 37°C, 98% humidity and 5% CO_2_ for 24 hours. All cell culture procedures were accomplished in a laminar flow hood, adhering to the sterility protocols of the materials and solutions.

### 0.7. Surgical Instruments & Adjunctives

Basic oral surgical instruments like pediatric extraction forceps, oral surgical kit, dental explorers, Metzenbaum scissors, towel clamps were obtained from GDC instruments, India. Hemostats, tissue pliers were purchased from Hu-Friedy, Chicago, IL, USA. Mallet hammer, Hohman’s bone spike, bone spike A.O type, Farabeuf periosteum elevator were acquired from Uma Surgicals, Mumbai, India. PeroxiDam, a light-curing material was purchased from Latus, Ukraine. Additionally, flexible blades were fabricated by flattening the syringe needles (3/8 inch - 3 1/2 inches, BD Precision Glide™ needles) in the workshop of MOD Lab, BARC, Mumbai for severing Sharpey’s fibers within the periodontal ligament. The blades were sharpened in the grinding motor by an extremely delicate touch for obtaining sharp contour at the edges^53^ by an experienced bioengineer and surgeon. All surgical instruments and accessories were autoclaved following standard protocols.^54^

### 0.8. Experimental Surgical Protocol For Customized Dental Implants Placed In Mandible Of Rabbits

For clinical chemistry, hemogram and other relevant determinants, the sampling of blood (*∼* 3 ml) from rabbits were performed prior to the day of surgery (Supplementary Tables). Pre-operative assessment (extraoral and intraoral) along with administration of Enrofloxacin (Bayrocin, Bayer) intramuscularly (5 mg kg^−1^) q.d., was continued for 3 days. On the day of surgery, induction of surgical anaesthesia was achieved in rabbits with intramuscular injection of 35 mg kg^−1^body weight of ketamine and 5 mg kg^−1^body weight of xylazine (Indian Immunologicals). The intraoral region was cleaned prior to surgery with antiseptic 2% chlorhexidine gluconate and 2% povidone-iodine solution. The experimental surgery was performed by an experienced bioengineer and surgeon under the supervision of a senior scientific officer at RBHSD, BARC, Mumbai, India. Apart from the usual oral surgical instruments, innovative and delicate surgical instruments such as the aforementioned modified thin flexible blades, specific to the experimental surgical protocol were used for extracting the tooth, atraumatically. Briefly, the sterile modified blades were placed into the marginal gingiva of 301 in rabbits with the tip of the blade proceeding apically whilst applying gentle finger pressure on the exterior of attached gingiva, thereby guiding the blade in a careful and delicate oscillatory (forward and backward) linear motion around the cervical^3*rd*^of tooth circumference for tearing the gingival fibres. Later, an extended flexible blade following closely the curvature of 301 in rabbits was gently inserted (towards the extremity i.e. apically involving middle^3*rd*^ and apical^3*rd*^ of the tooth surface) under sterile saline-solution irrigation, during which the tip of the index finger of the opposite hand was placed on the corresponding attached gingiva to achieve stability and accurately placing the blade in the periodontal ligament space. The blade within the periodontal space is gently twisted with minimal pressure and secured for 20-30 secs for setting free the periodontal attachment fibres. Thereafter, the blade was further explored in the periodontal space, moving all the way around the tooth with identical surgical manoeuvre. The loose tooth, 301 is gently grasped by a pediatric extraction forceps and pulled out of the alveolar socket.

**FIG. 2.**
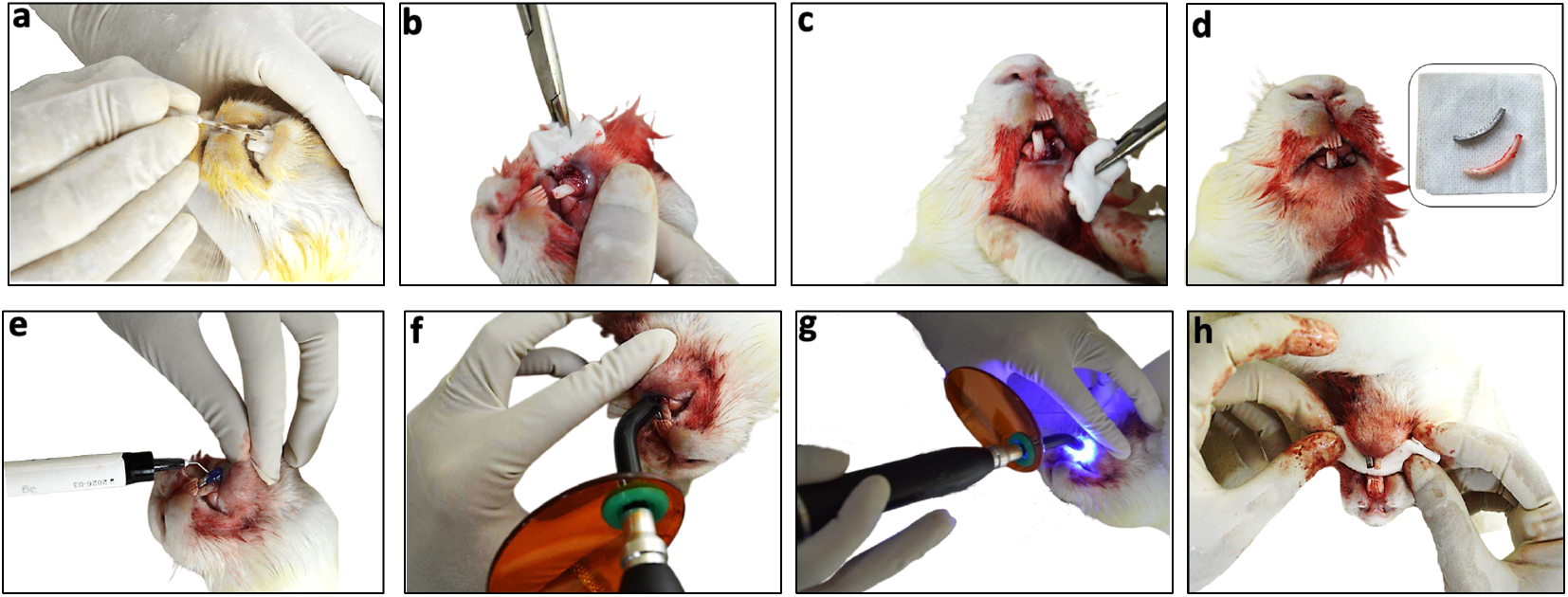
Representative intraoperative photographs revealing the experimental surgical technique for installing titanium dental implant in the jaw of rabbit study models for achieving the desired functionality of “proprioception” in implants: **(a)** gentle insertion of the customized blade (flexible enough to follow the curvature of the entire root, preferably dimension of the blade should be smaller than that of the root) into the gingival sulcus and later continuing deep into the periodontal space with the tip of the blade angled towards the long axis of the tooth followed by cautious rocking motion around the entire tooth circumference for disengaging gingival and periodontal fibres respectively. The described surgical manoeuvre leads to severance of connective tissue attachment fibers bridging the tooth and alveolar bone and will render the tooth mobile and loose. Root fractures are avoided carefully. (b-c) inspection of the surgical site after grasping the tooth with the pediatric extraction forceps and applying gentle twisting / semi-rotatory motion along the transverse axis, together with the careful pulling of the tooth from the socket. (d) image of the oro-facial region of the rabbit study model after the extraction of mandibular left central incisor with an inset image showing a 3D-printed titanium dental implant i.e. the replacement model of the mandibular left central incisor and the actual extracted mandibular left central incisor tooth. Because coated test implant being white does not provide sharp coherent and intelligible information, an uncoated titanium implant was chosen for representing a good contrast in the intraoperative photographs. The primary stability of the implant is achieved by placing the sterile media wetted, push-fit implant carefully conforming its long axis to that in the entrance of the socket and later gently tapping it in the empty socket while stabilizing and supporting the jaw carefully. Dimensions of implant and tooth in all planes along with its curvature, coating thickness etc. needs to be accurately estimated to avoid complications like splitting of the socket during insertion of an implant which may further lead to complications/sequelae e.g. submental space infection in relation to mandibular anteriors etc.(e) after appreciative biomechanical stability of dental implant was achieved in the mandible of rabbit study models, a light-curing material based on methacrylate resin, was applied to protect the wound during the initial healing phase.The application area was 2 - 3.5 mm wide and approximately 1.5 - 1.8 mm thick.The material should adhere excellently and provide an impervious seal for both types of interfaces: peri-mucosal and endosseous interfaces so as to decrease the risk of surgical site infection, bleeding and negating the possibility of saliva contamination in the operative site. The application started covering the facial and lingual aspects of the bone-implant interface followed by using slight pressure proximally directed to the inter-dental space on either side.(f-g) the light-curing material was cured for 20 - 30 seconds using a polymerisation lamp, evenly in a scanning motion.(h) post-surgical inspection of the installed titanium implant, soft tissue and adjacent anatomical structures etc.

Proper installation of the customized endosseous dental implant after simple and clean surgical extraction of 301 is the most crucial step. The primary stability of the implant (length = 2 mm shorter than the total length of respective 301) is achieved by placing the push-fit implant carefully in the fresh extraction socket and later gently tapping it with an orthopaedic mallet for its advancement in the entire length of the fresh empty socket. Before placement, the implant is wetted with sterile culture media for its easy gliding. The topmost, outer surface of the implant to be tapped should not be bevelled. Rather it should be flat for even contact of mallet surface (during gentle tapping) resulting in proper distribution of forces and uniform advancement of the implant along its long axis. Dimensions of implant and respective tooth i.e. 301 along with its curvature is calculated with great precision (Supplementary Fig) to avoid complications like splitting of the socket during insertion of implant etc. During the push-fitting of implant the mandible of rabbit is stabilized and supported carefully. A light-curing resin based barrier (PeroxiDam, Latus, Ukraine) was used to secure the interface of peri-implant soft tissue and titanium implants from rest of the oral cavity. Additionally, rabbit no. 3 with test implant received intramuscular 500 *l* q.d. of neurobion forte (Procter & Gamble Health Ltd. India) as a peripheral nerve recovery agent. Post-operative pain control was achieved with subcutaneous administration of 0.05 mg kg^−1^ buprenorphine hydrochloride. The rabbits were placed in individual cages during recovery. Clinical assessment, detection of possible radiographic changes from routine postoperative radiographs, physical examinations, monitoring etc. were included in the post-operative care and management. After euthanasia, histopathological patterns of superficial cervical lymph nodes (Supplementary Fig), peri-implant mucosa, tissue samples from spleen, liver, lung and kidney were evaluated. The relevant images during surgery were recorded and captured by a Nikon B500 Camera.

## Results Of Experimental Surgery

Essential findings of this proof-of-concept trial and its surgical outcome were performed at RBHSD, BARC, (Mumbai, India) and provides conclusive evidence, indicating towards one of the advanced possibility of biodegradable nanofibres coated implants. All surgeries were performed under general anesthesia by an experienced bioengineer and surgeon. Immediate postoperative examination revealed excellent primary stability for all implants (test and controls) placed in the mandible of all rabbit study models. The post-operative assessment was carried out at the end of 24 and 48 hrs. Further examination of the peri-implant site in the following days revealed normal and smooth tissue contour without any evidence of discoloration, exudation, or crusting. All the rabbit study models were found to constantly groom themselves. Subcutaneous administration of 0.05 mg kg^−1^ buprenorphine hydrochloride for all rabbits and 500 μl q.d. of neurobion forte i.m. for rabbit no. 3 with test implants, additionally, were continued. A sterile and soft diet with clean water was provided during the recovery period. Assisted feeding was not attempted, as the rabbit(s) started eating by itself within 24-36 hrs, post-surgery. It was observed that the rabbits were using the prehensile lips to assist in grasping softened food, which was later masticated by the combined action of the tongue and posterior teeth. However, at the beginning of the 8^*th*^ day, upon close examination, the implants were noted to be displaced along the longitudinal axis and were slightly shifted, mostly in a medial and posterior direction. Examination of implants in the subsequent days confirmed various grades of mobility. Additionally, significant peri-implant probing depth on mesial and distal aspects for both test and control implants were noted in all rabbit study models when compared to that of a normal tooth. We found all implants had some measurable grades of mobility at their respective interface; however, we do not have strong evidence to suggest the specific and clear reason for such adverse effect on the bio-integration of test and control implants in spite of such implants achieving initial firmness in bone with excellent primary stability during surgery. Nevertheless, these findings suggest that implants of such dimensions are at risk of exposure to some level of opposing occlusal forces thereby transferring and directing those to the interface between the alveolar bone and implant. In all likelihood, the opposing structures mainly the corresponding maxillary tooth occluding the dental implant resulted in undesirable changes in the peri-implant bone level around the dental implant, leading to its loosening and subsequent dislodgement. It is noted that, after immediate extraction, the selected dental implants replacing the tooth shouldn’t protrude more than 0.2 - 0.5 mm from the alveolar ridge to avert such occurrence. Following a thorough evaluation and taking into account the extent of progressive bone loss and the effect of possible surgical intervention with newer sets of implants (test and controls) with smaller dimensions replacing failed ones at the exact surgical site, were planned. However, remnants of normal physiological and healthy periodontal apparatus are necessary to accomplish the objectives successfully. Hence, a decision was made to carry forward the research work with the exact surgical procedures described here, in another set of animal study models with smaller dimensions of test and control implants.

## Discussion

Restoration of malfunctioned or damaged peripheral nervous systems is an enormous challenge for the therapeutic paradigms of stem cells. Osseointegrated dental implants are associated with the disruption of proprioceptive pathways of teeth due to the missing PDL. Apart from the prosthodontic treatment option serve by a dental implant, neural regeneration, and repair at its interface are additional research objectives that have been taken in recently, towards the effective response, together as an *“advanced and improved prosthetic remedy”* required for complete physiological recovery of the masticatory apparatus.

The present proof-of-concept trial deals with targeted repairing of the nerve terminals that are present as an extension of the trigeminal system at the interface of the immediately placed dental implant and alveolar bone, thereby completing the neural circuitry of a dental proprioceptive route from the interface of a dental implant to the mesencephalic nucleus in the midbrain. Thus, in addition to the endogenous stem cells in the peri-implant region, exogenous mesenchymal stem cells seeded in the orthotopically placed nanofibrous coated implant in rabbit’s mandible, was utilized for neurogenesis/neural repair in peri-implant tissues so as to later gauge its proprioceptive features.

The results of our experimental surgery described herein include the actual extraction of the complete tooth and installing an implant with minimal trauma to the adjacent structures. It is noted in the trial that using force without proper detachment of fibres would most likely induce a transverse fracture of the tooth at or below the gingival sulcus, making extraction with forceps impossible. Therefore, the described surgical technique requires thorough training for the knowledge and understanding of the related anatomical structures and implementation of the precise surgical skill, for executing the installation strategies to induce proprioceptive features in dental implants.

The PCL/gelatine nanofibers, such as ours, supports the nerve cells and improve the neurite outgrowth and cell differentiation process.^55^ Additionally, it has been previously reported by us that the titanium implants coated with PCL/gelatine nanofibrous scaffold, supports DPSCs, their proliferation and subsequent neural differentiation.^42^To ensure that the exogenous stem cells precisely participate in the terminal nerve repair at the interface we have systemically administered *Neurobion Forte*, as a peripheral nerve recovery agent^56,57^ in the rabbit study model with test implant.

However, additional specific interventional measures such as the inclusion of growth factors in relation to peripheral nerve regeneration, such as Neurotrophin-3, Neurotrophin-4, Recombinant human beta-NGF, and Fibroblast growth factors (FGF) protein^58^ in the components of nanofibrous coating, could have certainly resulted in further improvement in the overall objectives of the trial.

Further, in the process of bio-integration of such nanofibrous coated implants, the coating will gradually deteriorate^59^ resulting in shifting of the implant-bone interface medially in relation to longitudinal axis of the dental implant. As the implantation period progresses, we assume that the available degradation site will most probably be occupied by tissues having features similar to periodontium, progressively repairing terminal nerve endings with anastomosis and finally bridging alveolar bone and implant surface.

Histological staining often permits initial recognition of the cellular and structural features of the regenerated tissue approximating the implant surfaces. The peculiarities of the repaired neural framework, secured in the confined periprosthetic tissue could be analysed by histological methods confirming morphological changes after nerve regeneration. Injuries such as those induced by exodontia may damage a significant population of the peripheral axons of sensory bipolar neurons innervating the teeth,^60^ initiating the mechanism of axonal regeneration at the injury site.^61^ Later, a high number of regenerating clusters with Schwann cells elaborate processes^62^ could be identified in the recovering region.

The restored trigeminal proprioceptive pathway together with its specialized sensory discriminative features could further be corroborated in preclinical animal models using electroencephalographic, magnetoencephalographic^63^ and fMRI approaches^64^ etc. (Supplementary Fig).

## Contributions

S.D. and J.B. designed the research plan, S.D. performed the experiments, surgery and other tasks related to the project, wrote the manuscript and made the figures. Results were reviewed by J.B, H.D.S. and S.D and modifications jointly done. S.D was jointly supervised by H.D.S. and J.B.

## Acknowledgments

The authors gratefully acknowledge IIT Bombay and BARC, Mumbai for access to Central and Departmental facilities for this largely self-funded work.The authors also wish to express their sincere thanks to BETiC facility for the 3D-printing of titanium implants.

## Funding

The work was jointly supported by IIT Bombay (*J*.*B. personal research funds*) and BARC, Mumbai, India.

## Notes

### Competing Interest Statement

The authors have declared no competing interest.

### Summary of Updates

Additional files

