## Supplemental Information for "Engineering Proprioceptive Implants via Surgical and Regenerative Approaches: Preliminary Interpretations"

Under peer review

### Contents

1. **Implants & Mandibular Incisor** ... **1**
2. **Atomic Force Microscopy (AFM) Analysis** ... **2**
3. **Lymph Node Histology Of Rabbit Study Models** ... **2**
4. **Hematology & Clinical Biochemistry Of Rabbit Study Models** ... **3**
5. **Weight Of Rabbit Study Models On The Day Of Surgery** ... **3**
6. **Radiographic Planning & Measurements In Rabbits** ... **4**
7. **Electroencephalographic (EEG) Recordings & Tooth Stimulation In Human** . **5**
8. **EEG From Brain Computer Interface (BCI) & Tooth Stimulation In Human** **6**
9. **Brain MRI & Trigeminal Mesencephalic Nucleus In Human** ... **7**
10. **Conclusion** ... **8**

**References** ... **8**

1
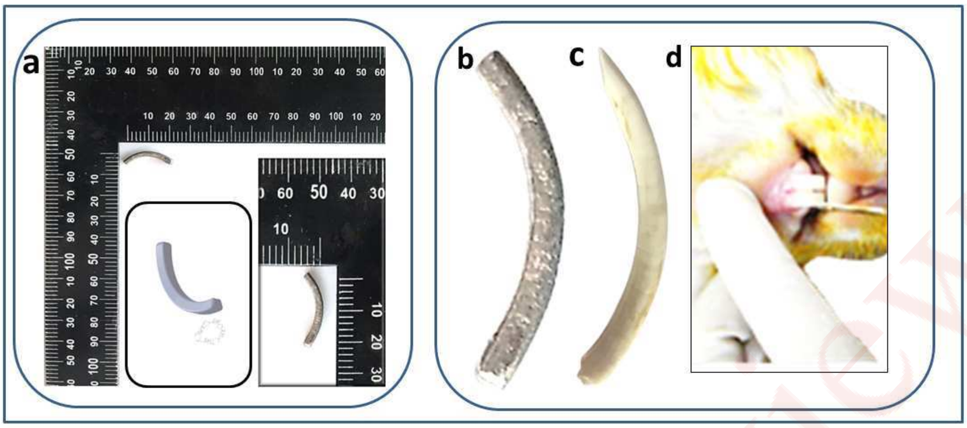


2
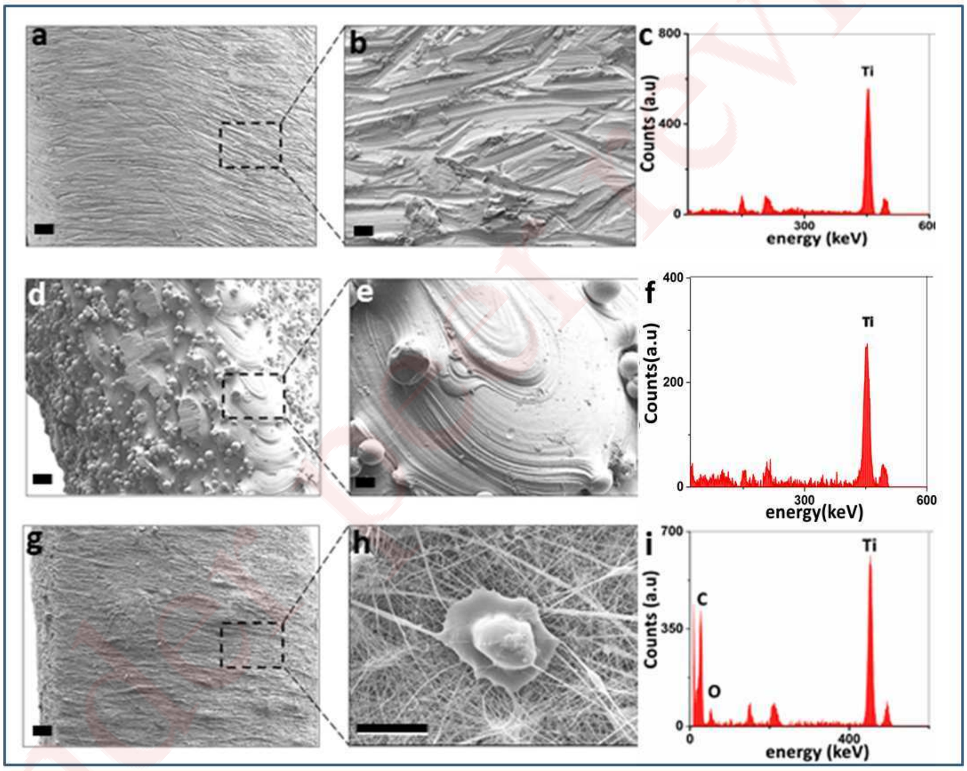


SF 1. **Implants and Mandibular Incisor:** (1). (a) Representative images of a 3D-printed titanium dental implant for rabbit's mandibular central incisor placed adjacent to a L-shape measuring tool with insets showing representative image of a design for the dental implant and its actual enlarge image placed adjacent to a measuring scale, (b) expanded image of a 3D-printed titanium dental implant, (c) extracted mandibular incisor from a rabbit study model (d) lower central incisor in the mandible of a rabbit just before surgical extraction. (2). Representative ESEM images and EDX analysis of control and test titanium dental implants, (a-b) manually fabricated titanium dental implant, (d-e) 3D-printed titanium dental implant, (g-h) titanium dental implant with nanofibers and undifferentiated dental pulp stem cells (DPSCs). Image scale bar represents: a, d and g - 100 µm; b, e -10 µm, and h – 20 µm. EDX of 3D-printed titanium dental implant (c), manually fabricated titanium dental implant (f), nanofibers coated titanium dental implant (i).The surface of manually fabricated titanium dental implant was roughened to promote osseointegration.

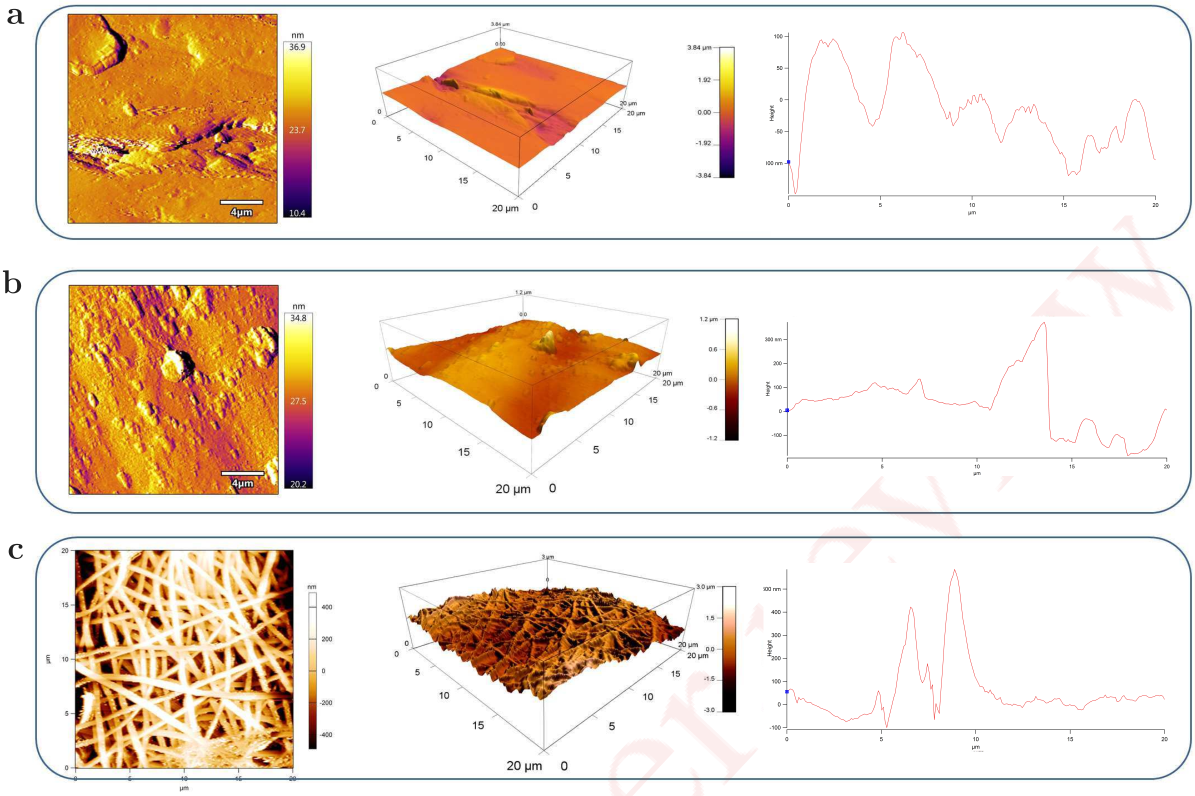


SF 2. **Atomic Force Microscopy (AFM) analysis:** AFM topographical images, three-dimensional images and profile plots in a 20 µm × 20 µm surface area for: (a) manually fabricated titanium dental implant (RMS surface roughness = 214.809 nm), (b) 3D-printed titanium dental implant (RMS surface roughness = 238.643 nm), (c) nanofibers coated titanium dental implant (RMS surface roughness = 468.627 nm) Control and test implant have different surface topographies as evident by the surface analysis. Nanofibres coated (Test) implants lead to an increase in the surface roughness when compared to those of 3D printed and manually fabricated titanium controls.

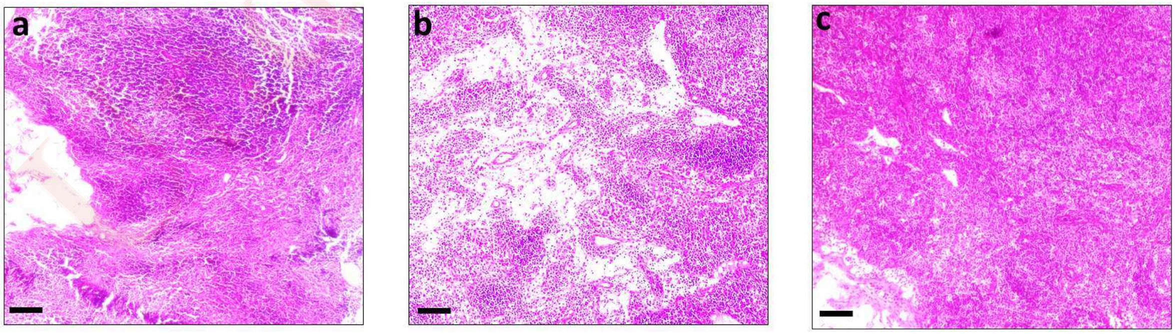


SF 3. **Lymph Node Histology:** Photomicrographs of superficial cervical lymph node for, (a) Rabbit 1 (Positive Control),* (b) Rabbit 2 (Positive Control)** and (c) Rabbit 3 (Test)*** study models. Histological evaluation showed no significant or obvious findings. Image scale bar represents: 100 µm. *Refer footnote in tables for classification of rabbit study models based on implant placement.*

TABLE 1 *Hematology and Clinical Biochemistry Of Rabbit Study Models*

| Parameters | Rabbit 1(Positive Control)* | Rabbit 2(Positive Control)** | Rabbit 3(Test)*** |
|---|---|---|---|
| Bili T | 0.5 | 0.3 | 0.2 |
| AST IU/L | 41 | 72 | 59 |
| ALT U/L | 42 | 56 | 71 |
| ALP IU/L | 147 | 201 | 193 |
| Total PRO g/dl | 7.1 | 6.9 | 7.2 |
| ALB g/dl | 4.2 | 3.9 | 4.1 |
| GLB g/dl | 2.9 | 3.0 | 3.1 |
| GGT U/L | 9.2 | 4.1 | 6.2 |
| Uric acid g/dl | 0.6 | 0.9 | 1.0 |
| BUN mg/dl | 29.1 | 20.8 | 31.3 |
| CREAT mg/dl | 1.5 | 1.4 | 1.5 |
| Na mEq/L | 138.9 | 139.5 | 141.3 |
| K mEq/L | 4.5 | 4.0 | 4.1 |
| Cl mEq/L | 98.2 | 103.4 | 104.1 |
| Ca mg/dl | 13.6 | 12.8 | 13.6 |
| P mg/dl | 5.1 | 4.9 | 3.9 |
| RBS mg/dl | 89 | 136 | 127 |
| Chol mg/dl | 81 | 48 | 61 |
| Tg mg/dl | 124 | 91 | 101 |
| Mg mg/dl | 2.3 | 3.3 | 2.8 |
| PT secs | 9 | 10 | 8 |
| APTT secs | 18 | 19 | 16 |

Source: Commercial laboratories
* Manually Fabricated Titanium Implant.
** 3D Printed Titanium Implant
*** Titanium Implant Coated with Nanofibres and Stem Cells

Under peer review

TABLE 2 *Weight Of Rabbit Study Models On The Day Of Surgery*

| Parameter | Rabbit 1(Positive Control)* | Rabbit 2(Positive Control)** | Rabbit 3(Test)*** |
|---|---|---|---|
| Weight (KGs) | 4.68 | 5.10 | 4.52 |

* Manually Fabricated Titanium Implant.
** 3D Printed Titanium Implant
*** Titanium Implant Coated with Nanofibres and Stem Cells

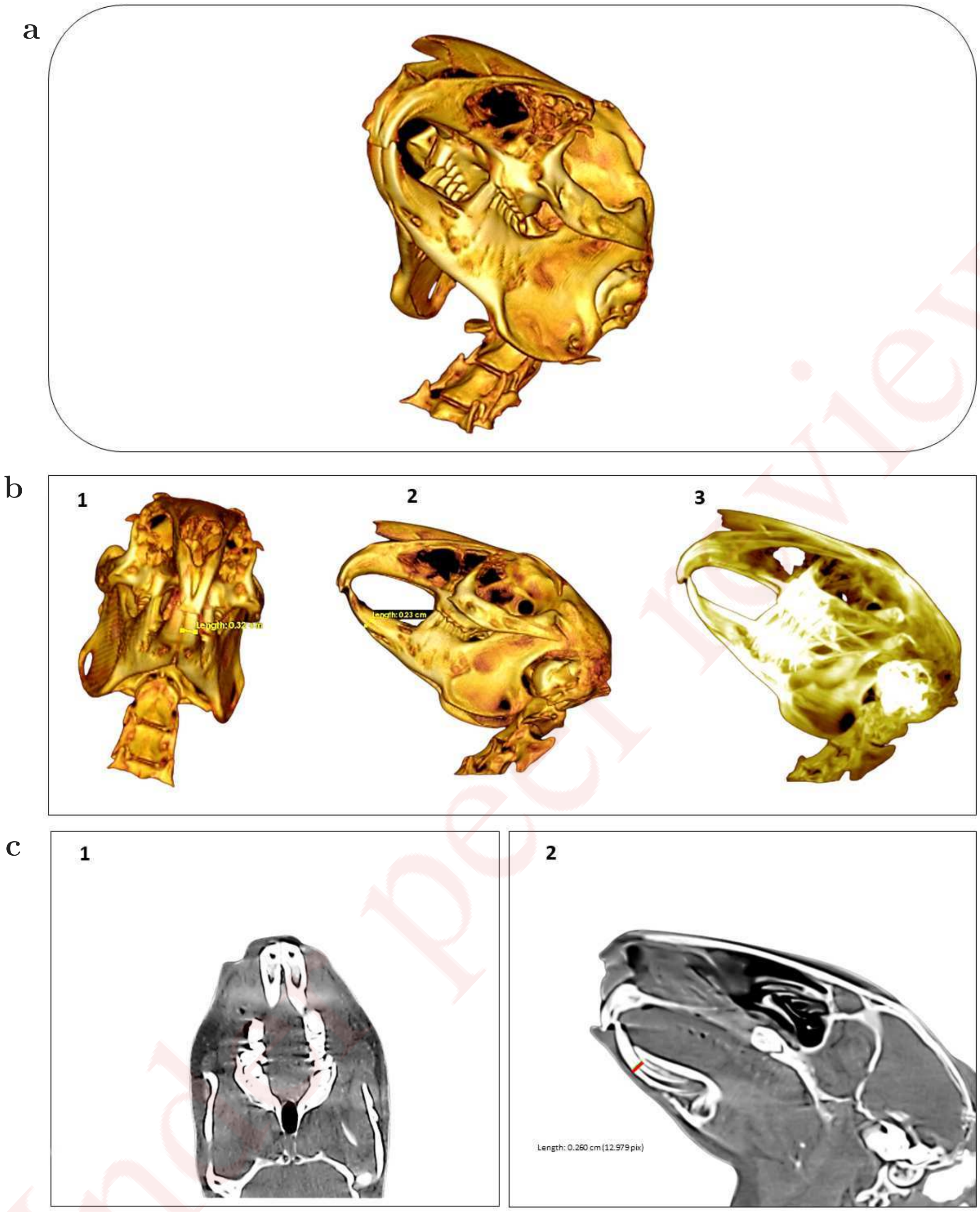


SF 4. **Radiographic Planning and Measurements:** Representative 2D and 3D CT reconstruction images for craniofacial region of a rabbit study model: (a) 3D-visualization of cranio-maxillofacial region before surgery, (b) measurements of mandibular central incisor to gauge its mesiodistal and buccolingual dimension in CT reconstruction models (1-2), 3D MIP of craniofacial region (3), (c) axial multislice CT (1) and sagittal multislice CT image (2), showing estimation of buccolingual length of mandibular incisor in a rabbit study model. Assessment of dentulous areas, interproximal bone height, counter of bone, the position, dimension and curvature of the tooth, periodontal space, anatomy and features of root, anatomic landmarks applicable for placing dental implants, corresponding tooth, opposing tooth and occlusion etc. were considered for surgical accuracy.

Under peer review

a

b

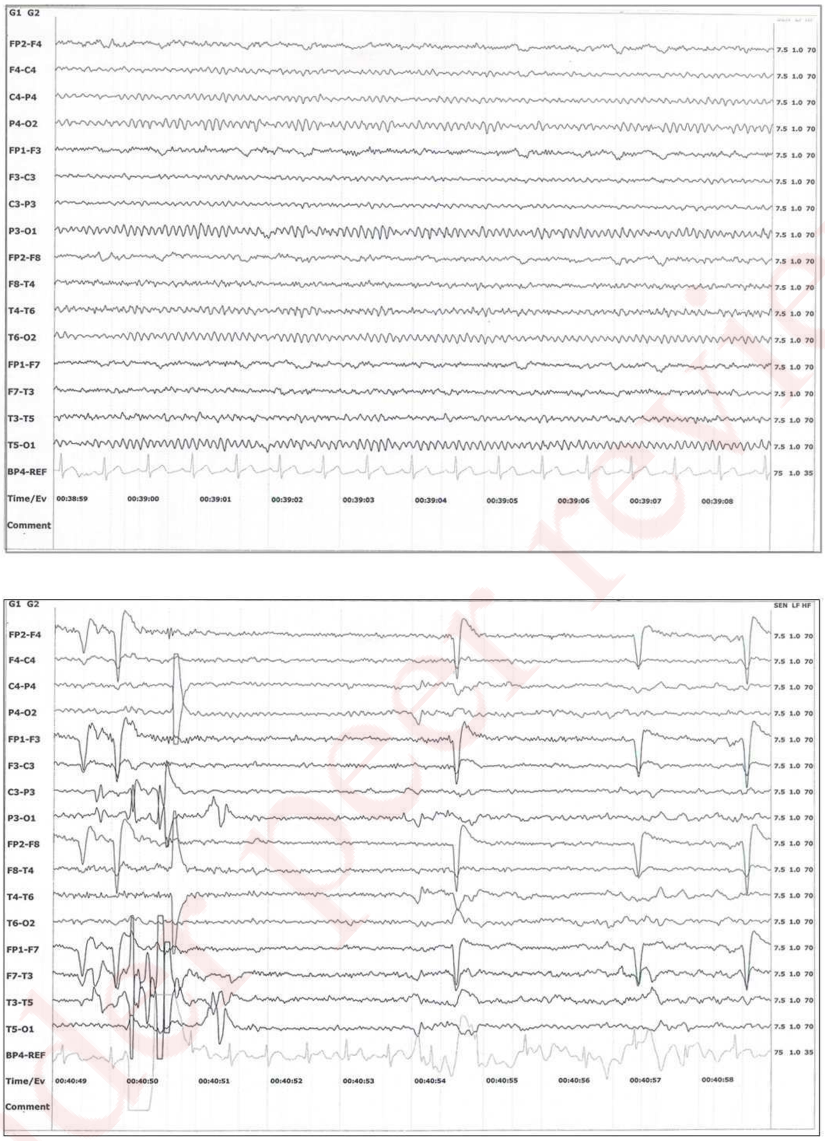


SF 5. **Electroencephalographic recordings in a 35 yrs. old healthy male subject with good dental health:** The international 10-20 system of electrode placement was used and the healthy subject lay supine during the EEG recordings (a) A fragment of EEG recorded during closed eyes. A normal 10 Hz Alpha rhythm is noted. (b) A fragment of EEG recorded during closed eyes in the same subject during teeth stimulation. The mechanical stimuli were gently delivered, perpendicularly to the longitudinal axis of the mandibular left central incisor tooth at about 1.5 - 2 Hz, through a bunt end probe. Physiological artifacts were noted in the electrodes placed nearer the eyes (FP1/2) and were mainly attributed to the eye ball movements. The trend of a downward deflection during the normal Bell's phenomena is observed. The out-of-phase deflection or phase-reversing is also noted in the electroencephalogram. No specific wave pattern in relation to tooth stimulus is detected. Stronger stimulus and refined data analysis of larger datasets might validate this method.

Under peer review

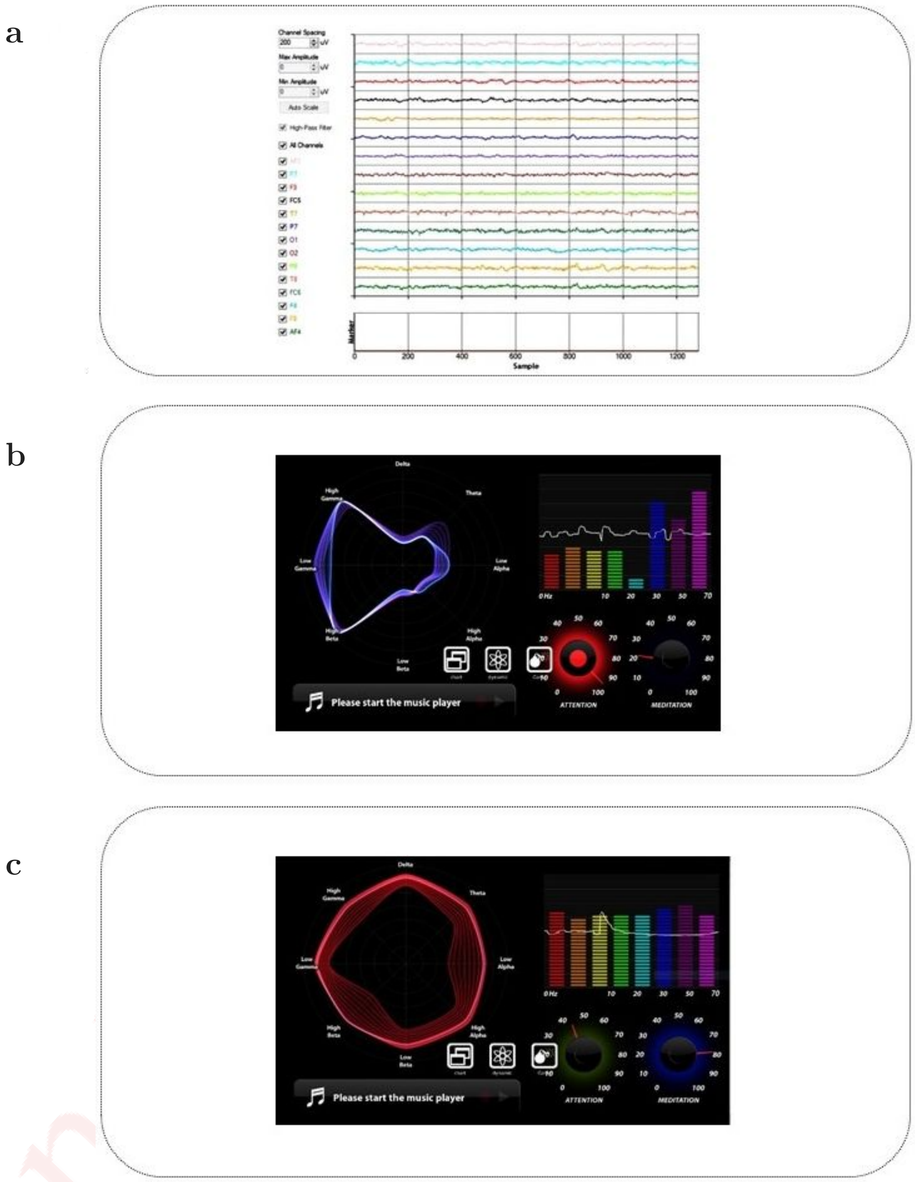


SF 6. **EEG From Brain Computer Interface (BCI) devices & Tooth Stimulation in a 35 yrs. old healthy male subject with good dental health:** Commercialized Brain-Computer Interface (BCI) devices such as (a) Emotiv EPOC headset (b) MindLink Brainwave sensor headband (c) NeuroSky MindWave headset that builds a wireless connection between the human brain and computers are used to sense EEG/Brainwave signals. However, these wearable EEG headset devices could not sense any specific electrical activity of the brain that could be correlated with tooth stimulation.

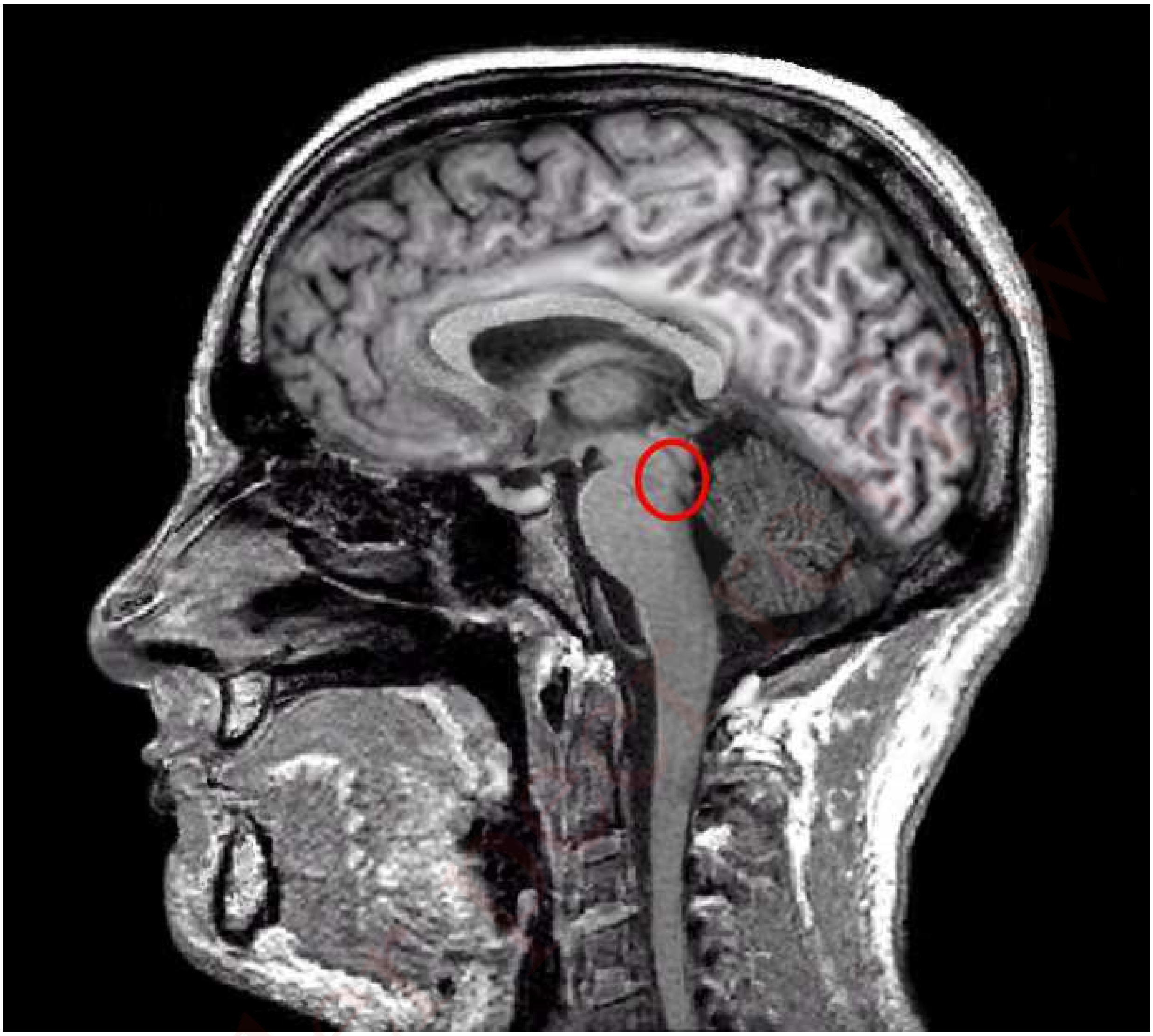

SF 7. **Brain MRI and Trigeminal Mesencephalic Nucleus in Human:** Sagittal magnetic resonance imaging (MRI) scan of a healthy, 35-year-old male showing the region of mesencephalic trigeminal nucleus located at the mesopontine junction (red circle) and is associated with the innervation of mechanoreceptors in the periodontal ligament. Very few investigations on the brainstem *fMRI* has been reported due to its correlation with numerous limitations such as technical challenges, complicated brain region, pulsatile signal from the cerebrospinal fluid, respiration induced brainstem movement, etc. Precise functional imaging such as BOLD signals through *fMRI* in the brainstem region are therefore much more demanding to detect than those in cortical areas.

### Conclusion

In the present proof-of-concept trial, an attempt has been made to restore the periodontium-like tissue around the dental implant, placed through immediate implantation in a fresh extraction socket in the mandible of a rabbit study model, thereafter, with a subsequent plan for gauging its proprioceptive features, after healing. Accordingly, the implant was coated with neuro-supportive nanofibers, seeded with undifferentiated mesenchymal stem cells, and placed in a rabbit study model, orthotopically. However, in spite of using smaller implants (i.e. *2 mm short than the total length of the actual tooth*), stresses were generated due to occlusal load resulting in a progressive bone loss in the peri-implant region for both test and control implants.Therefore, events related to implant loosening with peri-prosthetic bone loss and their subsequent dislodgement, were observed. Nevertheless, in the trial, the accuracy of the ***coating method***, (i.e. *with single or multiple motors placed between the syringe tip and collector plate in a modified electrospinning apparatus*) ***surgical protocol***, and ***primary stability*** of the implants during the experimental surgery were established. Advantageously, assisted feeding was not necessary and all rabbits survived the experimental surgery ( i.e. *minimal trauma were induced by the experimental surgery in the proof-of-concept trial*). It was also noted in the trial, that using shorter implants could have prevented the occurrence of peri-implant bone loss. Additionally, in such a developing scenario, the proprioceptive features of the regenerated periodontium-like tissue for test implants must be validated in the pre-clinical animal models before initiating clinical trials.

It has been established earlier that the stimulation of the periodontium results in a response in the maxillo-mandibular complex and simultaneously a reaction in the central nervous system, i.e. the higher mammals like humans (or *rabbits*) have a corresponding representation for the sensation of itself in the brain (*sensory homunculus*). Since we have placed our modified implant in rabbits, we have to depend on advanced non-invasive neuroimaging techniques such as *EEG*, *fMRI*, etc. for validation of the precise afferent feedback induced by the modified test implant(s). As the rabbit brain is not mapped extensively as the human brain, the *fMRI* response of a normal tooth, osseointegrated control implant and modified test implants for proprioceptive features placed in the rabbit jaw, could be compared to derive the conclusion. For providing stimulation, in an *fMRI* environment, a thread could be tied to the implant or tooth (i.e. *pulling of thread will provide the necessary stimulus*). Simultaneously, cortical projection, (*if any*) could be analyzed with normal tooth (*control or with standard sensory homunculus*), and inferences could be drawn. Likewise, *EEG* recordings from rabbits post-implant surgery, i.e. after the healing period could be performed. The extent of sensation could be evaluated based on a clinical scale from "0" (*no facial response*) to a series of higher numbers with respective well-defined facial expression responses based on the intensity of the mechanical stimulation on the implants or tooth.

Under peer review
